# Analyzing the covariance structure of plasma signaling proteins in relation to the diagnosis of dementia

**DOI:** 10.1101/2022.10.28.514320

**Authors:** Calvin Guan, Rhoda Au, Alvin Ang, Ashis Gangopadhyay

## Abstract

Numerous studies have shown that individuals with dementia have exhibited activation of inflammatory pathways in their brains. Typically, these studies use traditional and well-established regression methods for data analysis. In this paper, a new approach is introduced that utilizes the analysis of the covariance structure using methods related to the principal component analysis (PCA) theory. Eleven biomarkers related to neuroinflammation were used to determine the association with the onset of dementia. Various demographic covariates were adjusted to account for possible confounding effects of the covariance structure. Three hypothesis testing methods were considered to discern differences between partial covariance matrices for comparing power and Type I errors through simulation studies. Application of hypothesis testing methods using data from Framingham Heart Study (FHS) found significant differences in covariance matrices between the non-dementia and dementia groups.

## 1. Introduction

There is general research consensus that inflammation in pathologically vulnerable regions of the brain is associated with dementia, and specifically with the subtype of Alzheimer’s disease (AD; Morgan et al., 2019). Therefore, it has been suggested that treating inflammatory conditions could inhibit the development of dementia. Given that inflammatory molecules have been found in autopsy (Hopperton et al., 2018), it raises the question at what antemortem time point is neuroinflammation a risk factor for the onset of dementia later in life. Whether inflammation is a causal factor of dementia or simply a byproduct is still widely debated among the scientific and clinical communities (Wyss-Coray and Rogers, 2012). Epidemiological studies and treatment trials using anti-inflammatory drugs have been disappointing, although concerns have been raised about the methodological fidelity of the trials (Wyss-Coray and Rogers, 2012). As there is yet a proven and effective cure for dementia, it is crucial to understand the underlying mechanisms further to develop such treatments.

Protein biomarkers are used to quantify the level of inflammation in the brain. For this study, eleven measures that are recognized as inflammatory biomarkers are assessed as risk factors for dementia. The most common inflammatory biomarker is C-reactive protein (CRP), which is found in higher concentrations after an aneurysmal subarachnoid hemorrhage (SAH; Frontera et al., 2017). Other biomarkers include interleukin-6 (IL6) and osteoprotegerin (OPG). IL6 is a pleiotropic cytokine that contributes to host defense during infection and tissue injury (Kang et al., 2019), while OPG is a soluble secreted protein and decoy receptor that is associated with inflammation after ischemic stroke (Shimamura et al., 2014). These protein biomarkers are gathered by venous blood samples drawn from participants and then the proteins of interest are isolated, frozen, and measured (Staerk et al., 2020).

The Framingham Heart Study (FHS) has been collecting longitudinal data from three generations of participants for over 7 decades that includes demographics, inflammatory biomarkers, incident dementia and clinical co-morbidities. Traditional multi-variate statistical methods have been previously used to assess the relationship between inflammation and dementia risk, but these hypothesis testing procedures can produce false positives at an unacceptably high rate even with a few simultaneous tests. Principal component analysis (PCA) and partial covariance analysis offer an alternative approach. PCA approximates data by the product of the “object patterns” and the “variable patterns” (Wold et al., 1987). Although partial covariance adjusts the scale measures for a set of variables (Baba et al., 2004). These methods can summarize a set of variables and identify the fundamental structure of the set, which can be invaluable for understanding the underlying factors that affect the combined impact of these variables. A useful application of the group structure is to facilitate the comparison of a set of variables for different population groups.

For this study, three distinct methods for the hypothesis testing of equivalence of covariance matrices are applied to determine potential inflammation-related risk factors for dementia: Principal Component Group Comparison (Krzanowski, 1979), Parametric Bootstrap (Forkman et al., 2019), and the Tracy-Widom Statistic (Johnstone, 2009). Primary objectives are to compare the test statistics and p-values via various visualization tools, as well as to assess the Type I error rate and power of the hypothesis testing procedure via simulations. Given the novelty of the approaches utilized in this paper, it is imperative to perform simulation analysis for a comprehensive understanding of these inferential tools. For example, the simulation analysis allows visualizing and evaluating each method’s unique advantages and disadvantages within controlled settings. The PCA comparison method is intuitive, and the nonparametric framework requires fewer assumptions about the underlying data generating process but is untested in the context of hypothesis testing. The Tracy-Widom distribution is parametric; therefore, the p-values can be derived from the distribution by theoretical analysis. However, there are strict assumptions for the Tracy-Widom distribution, so the results may be questionable if the assumptions are not met. The Parametric Bootstrap method allows the computation of the p-values without the restrictive distributional assumptions. The results are further summarized into test statistics to test the differences in partial covariance matrices.

In summary, the paper discusses three novel approaches to understanding the differences between covariance structures of multivariate data and provides guidelines for using these methods using extensive simulation studies. The potential clinical contribution of these analyses is their application to confirm the role of inflammatory biomarkers as a risk factor for dementia.

## 2. Methodology and Materials

### 2.1. Design

In this section, we briefly describe the details of the study design.

#### 2.1.1. Participants

The Framingham Heart Study (FHS) is a community-based multi-generational prospective cohort study that began in 1948 to identify risk factors for cardiovascular disease and subsequently expanded to incorporate study of many common chronic diseases including dementia. For the current study, participants are members of the offspring cohort, which includes the biological children of the original FHS cohort and the spouses of the children (Feinleib et al., 1975). These participants had regular health examinations on average every four years. Data used for this analysis were collected at the seventh and eighth health examinations performed during 1998 - 2001 and 2005 – 2008, respectively (n = 2684).

#### 2.1.2. Inflammatory Biomarkers

Inflammatory biomarkers were measured from venous blood samples, as previously described (Staerk et al., 2020). Table 1 lists the 12 different biomarkers of inflammation (Fricker et al., 2019; Li et al., 2019).

**Table 1:**
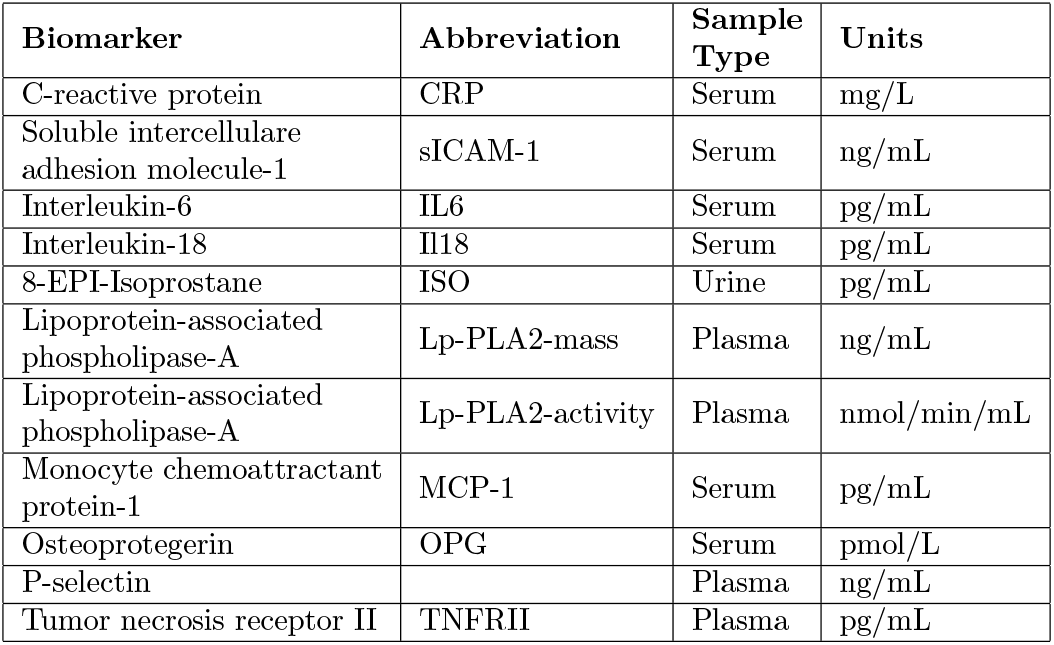
Biomarker Sampling Method

| Biomarker | Abbreviation | Sample Type | Units |
| --- | --- | --- | --- |
| C-reactive protein | CRP | Serum | mg/L |
| Soluble intercellular adhesion molecule-1 | sICAM-1 | Serum | ng/mL |
| Interleukin-6 | IL6 | Serum | pg/mL |
| Interleukin-18 | Il18 | Serum | pg/mL |
| 8-EPI-Isoprostane | ISO | Urine | pg/mL |
| Lipoprotein-associated phospholipase-A | Lp-PLA2-mass | Plasma | ng/mL |
| Lipoprotein-associated phospholipase-A | Lp-PLA2-activity | Plasma | nmol/min/mL |
| Monocyte chemoattractant protein-1 | MCP-1 | Serum | pg/mL |
| Osteoprotegerin | OPG | Serum | pmol/L |
| P-selectin |  | Plasma | ng/mL |
| Tumor necrosis receptor II | TNFRII | Plasma | pg/mL |

The biomarkers’ blood concentration levels are typically right-skewed, as there is a lower limit on the levels but no upper limit (Figure 1). Therefore, a logarithmic transformation is applied to biomarkers to reduce skewness and improve the performance of PCA decomposition.

**Figure 1:**
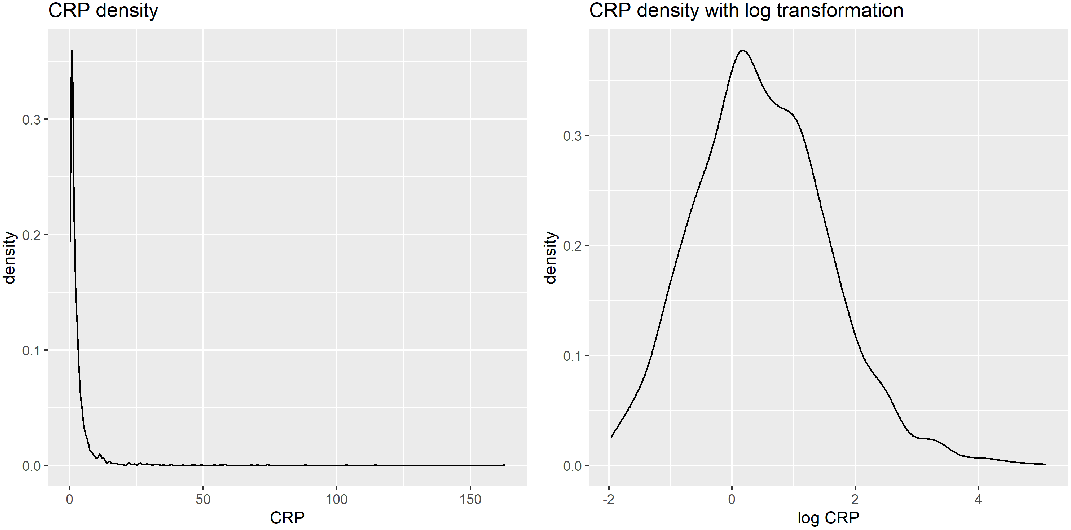
Kernel Density Estimation of CRP vs. Log-Transformed CRP

#### 2.1.3. Dementia Review

The dementia diagnosis of each participant in FHS is evaluated and verified through an adjudication panel that includes at least one neurologist and one neuropsychologist, who determined the diagnosis based on information from clinical examinations, medical records, and, when available, neuropsychological and neurological assessments and family interviews (Satizabal et al., 2016; Yuan et al., 2021). Ongoing surveillance for incident dementia that has been underway since 1976 identifies participants for diagnostic consideration. Therefore, those who do not undergo a dementia review assessment are presumed not to have dementia. Participants identified for possible mild cognitive impairment (MCI) were also excluded from the analysis (n = 158), to create a distinct separation between the dementia and non-dementia groups. Dementia is flagged whether the onset is before or after the biomarkers sampling.

#### 2.1.4. Adjusting for Covariates

Covariates that could significantly affect protein biomarker concentration are identified using rudimentary linear models and their mutual correlation. These covariates are age, sex, BMI, current smoking status, total cholesterol, ventricular rate, serum creatinine concentration and indicator of treatment for lipid disorders. Furthermore, the models are adjusted for the time difference between biomarker sampling and dementia diagnosis. Participants with missing covariates are excluded (n = 1021).

#### 2.1.5. Data Imputation

Missing data, be it due to non-attendance at the time of data collection, inconclusive results, unreadable notation, or something else, pose a significant challenge in a clinical setting. However, in traditional statistical analysis, a sample would be discarded if its variables had missing values. Instead, the PCA imputation method is applied on the logarithm of biomarker concentrations to preserve the sample size with minimal effect on the data quality. This method uses a regularized iterative PCA algorithm that fits an expectation-maximization (EM) algorithm to remove noise during the imputation step (Josse and Husson, 2012). Note that imputation works well when the proportion of missing values is not too high; as such, the inflammatory biomarker lysophosphatidic acid (lpa) was not considered in this analysis due to too many missing values (n = 1270).

### 2.2. Statistical Analyses

This section describes the various statistical techniques implemented to analyze the data along with a detailed discussion of the various characteristics of the methodologies.

#### 2.2.1. Participant Characteristics

After screening out exclusions and implementing imputation for missing observation, the data included 2684 participants, of which 2536 were in the non-dementia group and 148 in the dementia group. The study sample was characterized using means and percentages along with rudimentary univariate analysis to examine potential differences between the two groups. Table 2 provides the mean and standard deviation of continuous variables. The two-sample t-test of the mean difference between the dementia group and the non-dementia group is included for each variable. Table 3 - Table 5 show the counts and proportions for categorical variables, along with the *χ*^2^ test of independence. Age, BMI, creatinine concentration, total cholesterol, current smoking status, and lipid disorder treatment indicator show significant mean differences between the dementia and non-dementia groups whereas sex and ventricular rate were not significantly different. Interestingly, the dementia rate is slightly higher for females than males, which goes against the findings of contemporary studies (Podcasy and Epperson, 2016).

**Table 2:** Quantitative Covariates Descriptions

| Demographics | $\hat{\mu}_n$ | $\hat{\sigma}_n$ | $\hat{\mu}_d$ | $\hat{\sigma}_d$ | t | p - value |
| --- | --- | --- | --- | --- | --- | --- |
| Age | 65.285 | 8.516 | 77.453 | 6.193 | 22.681 | <0.001 |
| BMI (kg/m<sup>2</sup>) | 28.406 | 5.443 | 26.768 | 4.597 | -4.167 | <0.001 |
| Creatinine (mg/dL) | 0.908 | 0.299 | 0.986 | 0.332 | 2.817 | 0.005 |
| Total Cholesterol (mg/dL) | 186.432 | 37.576 | 179.115 | 35.855 | -2.407 | 0.017 |
| Ventricular Rate (beats / min.) | 62.336 | 10.415 | 63.101 | 11.380 | 0.799 | 0.426 |

$\hat{\mu}$ is the sample mean, $\hat{\sigma}$ is the sample standard deviation, **n** denotes the non-dementia group, and **d** the dementia group

**Table 3:** Covariates Variables Descriptions (Sex)

| Sex | N | N<sub>n</sub> | %<sub>n</sub> | N<sub>d</sub> | %<sub>d</sub> |
| --- | --- | --- | --- | --- | --- |
| Male | 1228 | 1164 | 0.459 | 64 | 0.432 |
| Female | 1456 | 1372 | 0.541 | 84 | 0.568 |

$\chi^2$ test of independence: $\chi^2 = 0.298$ , $p - value = 0.585$
**N** is the sample size of the respective group, while %<sub>n</sub> and %<sub>d</sub> are the proportion of the non-dementia and dementia group with given characteristics.

**Table 4:** Covariates Variables Descriptions (Current Smoking Status)

| Current Smoking Status | N | N<sub>n</sub> | %<sub>n</sub> | N<sub>d</sub> | %<sub>d</sub> |
| --- | --- | --- | --- | --- | --- |
| No | 2444 | 2302 | 0.908 | 142 | 0.959 |
| Yes | 240 | 234 | 0.092 | 6 | 0.041 |

$\chi^2$ test of independence: $\chi^2 = 3.983$ , $p - value = 0.046$

**Table 5:** Covariates Variables Descriptions (Treated for Lipids)

| Lipids | N | N<sub>n</sub> | %<sub>n</sub> | N<sub>d</sub> | %<sub>d</sub> |
| --- | --- | --- | --- | --- | --- |
| No | 1535 | 1476 | 0.582 | 59 | 0.399 |
| Yes | 1149 | 1060 | 0.418 | 89 | 0.601 |

$\chi^2$ test of independence: $\chi^2 = 18.464$ , $p - value < 0.001$

#### 2.2.2. Biomarker Characteristics

Table 6 provides the characterization of inflammatory biomarkers using the mean and standard deviation with two samples t-test of the mean difference as a comparison between the dementia and non-dementia groups. The results are mixed; four biomarkers have significant differences, while seven did not. However, this method evaluates biomarkers individually and is not able to detect potential inter-biomarker covariance differences. Furthermore, there may be demographic effects that cannot be captured using simple summary statistics. Therefore, it is necessary to use more sophisticated statistical methods to sufficiently understand the underlying covariance structure between the biomarkers.

**Table 6:** Log Predictor Descriptions

| Biomarker<br>(log concentration) | $\hat{\mu}_n$ | $\hat{\sigma}_n$ | $\hat{\mu}_d$ | $\hat{\sigma}_d$ | t | p – value |
| --- | --- | --- | --- | --- | --- | --- |
| CRP | 0.479 | 1.081 | 0.540 | 1.252 | 0.623 | 0.533 |
| sICAM-1 | 5.657 | 0.305 | 5.714 | 0.315 | 2.271 | 0.024 |
| IL6 | 0.622 | 0.728 | 0.993 | 0.773 | 6.031 | <0.001 |
| IL18 | 5.446 | 0.386 | 5.569 | 0.405 | 3.825 | <0.001 |
| ISO | 6.685 | 0.841 | 6.710 | 0.713 | 0.424 | 0.672 |
| Lp-PLA2-mass | 5.277 | 0.261 | 5.309 | 0.246 | 1.587 | 0.114 |
| Lp-PLA2-activity | 4.904 | 0.257 | 4.951 | 0.233 | 2.487 | 0.014 |
| MCP-1 | 5.901 | 0.307 | 6.014 | 0.341 | 4.181 | <0.001 |
| OPG | 1.532 | 0.302 | 1.816 | 0.326 | 10.952 | <0.001 |
| P-selectin | 3.666 | 0.318 | 3.687 | 0.327 | 0.771 | 0.442 |
| TNFR2 | 7.796 | 0.396 | 7.954 | 0.813 | 2.492 | 0.014 |

**Table 7:** Type I error and power under **DGP1**

| Method<sup>1</sup> | $n_0$ | $n_1$ | $\rho_1 = -.4$ | $\rho_1 = -.2$ | $\rho_1 = 0$ | $\rho_1 = .2$ | $\rho_1 = .4$ |
| --- | --- | --- | --- | --- | --- | --- | --- |
| PCGC | 500 | 500 | 0.8 | 0.61 | 0.052 | 0.63 | 0.816 |
| PCGC | 1000 | 1000 | 0.87 | 0.762 | 0.03 | 0.754 | 0.876 |
| PCGC | 1500 | 1500 | 0.892 | 0.804 | 0.044 | 0.79 | 0.89 |
| PBoot | 500 | 500 | 1 | 0.79 | 0.034 | 0.83 | 1 |
| PBoot | 1000 | 1000 | 1 | 0.986 | 0.048 | 0.98 | 1 |
| PBoot | 1500 | 1500 | 1 | 1 | 0.046 | 0.998 | 1 |
| TW | 500 | 500 | 0.992 | 0.334 | 0.004 | 0.342 | 0.994 |
| TW | 1000 | 1000 | 1 | 0.816 | 0.002 | 0.824 | 1 |
| TW | 1500 | 1500 | 1 | 0.98 | 0.004 | 0.974 | 1 |

$\rho_1 = 0$ constitutes the null distribution

**Table 8:** Type I error and power under **DGP2**

| Method<sup>1</sup> | $n_0$ | $n_1$ | p | $\theta_1 = 1$ | $\theta_1 = 2$ | $\theta_1 = 3$ | $\theta_1 = 4$ | $\theta_1 = 5$ |
| --- | --- | --- | --- | --- | --- | --- | --- | --- |
| PCGC | 500 | 500 | 10 | 0.05 | 0.056 | 0.052 | 0.05 | 0.06 |
| PCGC | 1000 | 1000 | 10 | 0.046 | 0.046 | 0.046 | 0.046 | 0.042 |
| PCGC | 1500 | 1500 | 10 | 0.056 | 0.046 | 0.044 | 0.048 | 0.05 |
| PBoot | 500 | 500 | 10 | 1 | 1 | 0.05 | 0.762 | 1 |
| PBoot | 1000 | 1000 | 10 | 1 | 1 | 0.074 | 1 | 1 |
| PBoot | 1500 | 1500 | 10 | 1 | 1 | 0.064 | 1 | 1 |
| TW | 500 | 500 | 10 | 0.074 | 0.008 | 0.046 | 1 | 1 |
| TW | 1000 | 1000 | 10 | 0.192 | 0.026 | 0.062 | 1 | 1 |
| TW | 1500 | 1500 | 10 | 0.33 | 0.036 | 0.052 | 1 | 1 |

$\theta_1 = 3$ constitutes the null distribution

**Table 9:**
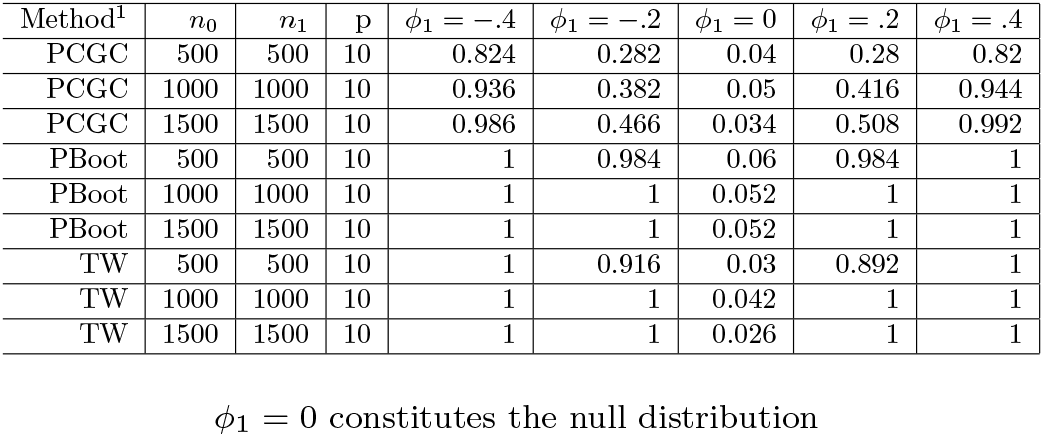
Type I error and Power under **DGP3**

| Method<sup>1</sup> | $n_0$ | $n_1$ | p | $\phi_1 = -.4$ | $\phi_1 = -.2$ | $\phi_1 = 0$ | $\phi_1 = .2$ | $\phi_1 = .4$ |
| --- | --- | --- | --- | --- | --- | --- | --- | --- |
| PCGC | 500 | 500 | 10 | 0.824 | 0.282 | 0.04 | 0.28 | 0.82 |
| PCGC | 1000 | 1000 | 10 | 0.936 | 0.382 | 0.05 | 0.416 | 0.944 |
| PCGC | 1500 | 1500 | 10 | 0.986 | 0.466 | 0.034 | 0.508 | 0.992 |
| PBoot | 500 | 500 | 10 | 1 | 0.984 | 0.06 | 0.984 | 1 |
| PBoot | 1000 | 1000 | 10 | 1 | 1 | 0.052 | 1 | 1 |
| PBoot | 1500 | 1500 | 10 | 1 | 1 | 0.052 | 1 | 1 |
| TW | 500 | 500 | 10 | 1 | 0.916 | 0.03 | 0.892 | 1 |
| TW | 1000 | 1000 | 10 | 1 | 1 | 0.042 | 1 | 1 |
| TW | 1500 | 1500 | 10 | 1 | 1 | 0.026 | 1 | 1 |

$\phi_1 = 0$ constitutes the null distribution

**Table 10:** Data Analysis, Test Statistics and P-values

|  | Simple Covariance | Partial Covariance |
| --- | --- | --- |
| PCGC | 0.038 (0.64) | 1.531 (0.01) |
| PBoot | 0.947 (0) | 0.953 (0) |
| Tracy-Widom | 86.333 (0) | 91.248 (0) |

#### 2.2.3. Partial Covariance

A multivariate data covariance matrix measures the joint structural association among a set of variables. Comparison of covariance matrices is a key tool for understanding differences between multivariate structures of multiple groups of data. This approach has been extensively used in the study of a wide range of problems involving multidimensional data, such as understanding the role of genetic constraints in the determination of evolutionary trajectories in adaptive radiation (Colautti and Barrett, 2011), the response of the genetic architecture to environmental heterogeneity (Robinson et al., 2009), the evolution of phenotypic integration (Kolbe et al., 2011), (Monteiro and Nogueira, 2010), multicharacter phenotypic plasticity (Mallitt et al., 2010) and sexual dimorphism (Barker et al., 2010) (Campbell et al., 2011), among others.

The partial covariance allows exploration of the structural association between the variables of interest after adjusting for exogenous variables. The logarithmic transformed biomarkers are adjusted by controlling for covariates. This is done by fitting a linear model for each biomarker against the covariates and estimating the covariance using the resulting residuals. Therefore, data analysis uses partial covariance matrices instead of traditional covariance matrices to account for demographic effects.

#### 2.2.4. Preliminary Screening for the Association of Inflammatory Biomarkers with Dementia

To explore the relationship of inflammatory biomarkers with dementia status, the partial covariance structure of biomarkers is compared between the two groups. Visual comparisons of the partial correlation matrices between the two groups were made using heatmaps and scree plots. The heatmap visualizes high values using reddish colors and low values using blueish colors, akin to a temperature map. A scree plot is a graphical tool that plots the eigenvalues, i.e. amount of variance explained by each principal component, of the covariance matrix by decreasing orders of magnitude. The scree plot can be used to find the number of significant components to keep in a principal component decomposition or the number of dominant factors in factor analysis (Cattell, 1966). These descriptive tools were utilized to identify potential structural differences between the partial covariance matrices of the non-dementia and dementia groups.

#### 2.2.5. Hypothesis Testing

Hypothesis testing methods served as a reference analytical tool to assess the difference between the partial covariance matrices of the two groups.

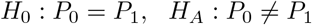

where *P*_0_ and *P*_1_ are the population partial covariance matrices of the non-dementia and dementia groups, respectively. A hypothesis testing method calculates a test statistic and the corresponding p-value from the data that are used to make a decision. Three novel approaches to the hypothesis testing problem were considered, namely, Principal Component Group Comparisons, Parametric Bootstrap, and Tracy-Widom statistic.

#### 2.2.6. Principal Components Group Comparison

The first method to assess the differences between two covariance matrices was the Principal Components Group Comparison (PCGC) method based on the comparison of principal components of different groups (Krzanowski, 1979). Let *L* and *M* be the principal component loading matrices of the non-dementia and dementia groups, respectively, then the test statistic is the minimum angle between the space of the first component, given by

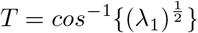

where *λ*_1_ is the first eigenvalue of *S* = *LM* ′*ML* ′.

A small minimum angle implies that the two groups are similar with respect to the first principal component. As the minimum angle does not follow a known probability distribution, a bootstrap method is used to calculate the p-value. Bootstrap method creates *B* bootstrap data by randomly shuffling the group labels. Then, a bootstrap test statistic *T*_*b*_ is calculated for each bootstrap data (*b* = 1, …, *B*). Finally, the p-value is calculated as the proportion of bootstrap samples in which the observed test statistic is smaller than the bootstrap test statistics.

#### 2.2.7. Parametric Bootstrap

The Parametric Bootstrap test takes inspiration from (Forkman et al., 2019), where the authors propose a hypothesis testing method for the number of significant principal components in a standardized data matrix. First, the authors simulate simple parametric bootstrap samples and calculate the test statistic for each sample. The bootstrap is parametric because it assumes a normal distribution. It is relatively simple approach because it is based on a standard distribution with no estimation of parameters. Then, the bootstrap test statistics are compared to the observed test statistic to calculate the bootstrap p-value. Estimation of parameters are not needed because the observed data are already standardized to zero mean and unit variance.

The methods in Forkman et al. (2019) are then modified to test the hypothesis of the difference between two covariance matrices. First the test statistic is calculated as such:

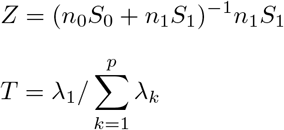

Where *S*_0_ and *S*_1_ are the sample partial covariance matrices of the standardized data from the dementia group and the non-dementia group, and *n*_0_ and *n*_1_ are the respective sample sizes. Furthermore, *T* is the test statistic and *λ*_*k*_ is the *k*th eigenvalue of *Z*.

In summary, the test statistic is calculated by combining the covariance matrices of two groups in such a way that if the two matrices are equal, the largest eigenvalue of the combined matrices follows the greatest root statistic distribution (Johnstone, 2009).

The p-value is then calculated using the following bootstrap algorithm:

##### Algorithm 1

Parametric Bootstrap P-Value

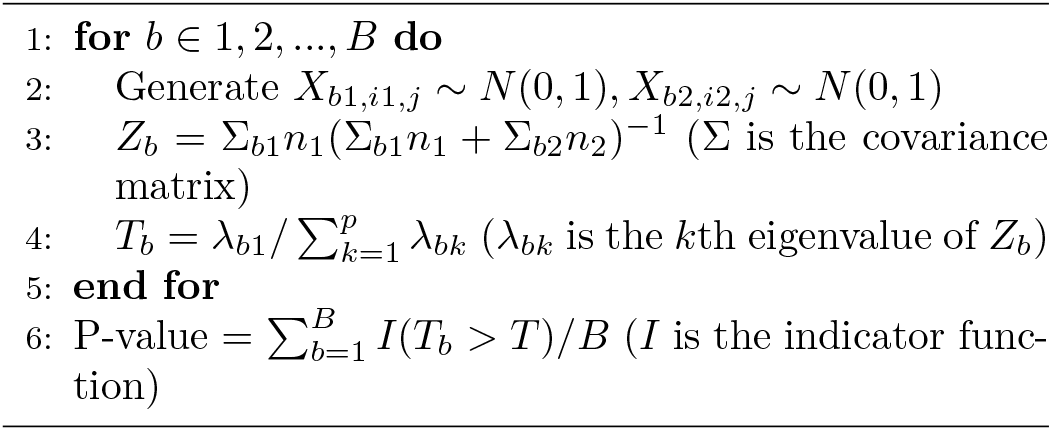

#### 2.2.8. Tracy-Widom Statistic

The Tracy-Widom distribution was introduced in (Tracy and Widom, 1994), as the probability distribution of the normalized eigenvalue of a random Hermitian matrix. Johnstone (2001) established the use of the Tracy-Widom distribution of order 1 as the asymptotic distribution of the largest eigenvalue in the covariance matrix of independent Gaussian variables. A follow-up paper (Johnstone, 2009) explores the application of the Tracy-Widom distribution of order 1 to multivariate analysis, of relevance is the group comparisons of covariance matrices.

The hypothesis tested considered *S*_0_ and *S*_1_, the sample covariance matrices for both groups. Let *n*_0_ be the number of participants in the non-dementia group, and *n*_1_ for the dementia group. And let *λ*_1_ be the largest eigenvalue of *S* = (*n*_0_*S*_0_ + *n*_1_*S*_1_)^−1^*n*_1_*S*_1_. Finally, define *F*_1_ as the Tracy-Widom distribution of order 1 and *p* the number of predictors, then under *H*_0_:

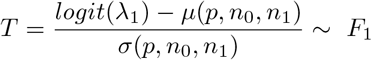

Where *µ* is the centering term, and *σ* the scaling term, defined as:

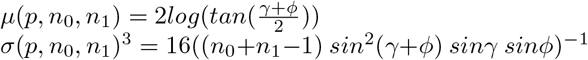

where

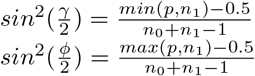

Therefore, *T* is the test statistic, and the p-value is given by *Pr*(*T > t*|*H*_0_) = 1 − *F* (*t*).

In practice, we approximate the Tracy-Widom p-value using a Gamma distribution as outlined by (Chiani, 2014), as the Tracy-Widom p-value is difficult to calculate directly.

#### 2.2.9. Simulation Study

Before addressing the core question related to differences in inflammatory biomarkers between the non-dementia group and the dementia group, it is important to first compare the performances of the three hypothesis testing methods discussed in the last section using simulation studies. Simulation is necessary to evaluate and compare the results of the three methods in a controlled setting, as established theoretical results for these methods are limited. Two groups of random variables are generated from a parametric distribution in the simulations. The first group had a fixed parameter, while the second group varied the parameter in a range where the fixed parameter of the first group is the center. The goal of these simulations is to visualize the change in statistic and p-values as the distributions of the two groups differ. The Type I error and power of these tests are assessed below. The following describes the data-generating process (DGP) for each simulation below.

##### DGP1

Data is generated from bivariate normal distributions with zero-mean and unit variance, **X**_**0**_, **X**_**1**_, where the correlation is defined as *ρ*_0_ = 0 *ρ*_1_ ∈ (−1, 1).

##### DGP2

Following the work of (Zheng et al., 2017), let **x**_**ki**_ = (*x*_*k*1*i*_, …, *x*_*kpi*_) represent the predictor vector for the *i*th observation in group *k*, in this case *k* = 0, 1 and generate *x*_*kji*_ = *w*_*kji*_ +2*w*_*k,j*+1,*i*_ +*θ*_*k*_*w*_*k,j*+2,*i*_, where *w*_*kji*_ ∼*N* (0, 1). Parameters are defined as *θ*_0_ = 3, *θ*_1_ ∈ (1, 5). This simulation creates a complex dependency based on a moving average process that is not directly obvious.

##### DGP3

Designed as a multivariate normal simulation with *X*_*k*_ ∼ *N* (0, Σ_*k*_) *k* = 0, 1. Where the *i* th row and *j* th column of Σ_*k*_, *σ*_*ijk*_, is defined as 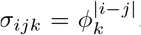, which follows the correlation structure of an autoregressive process of order 1 (AR(1)). Specifically, the parameters are defined as *φ*_0_ = .5, *φ*_1_ ∈ (0, 1). This simulation can be described as a multivariate variation of **DGP1**.

In each simulation, the test statistic and the p-value are graphically visualized. A fine grid of discrete values is set for *ρ*_1_, *θ*_1_, and *φ*_1_ respectively. Each DGP is simulated 10 times, and the average of each set of test statistic or p-values against their respective value of *ρ*_1_, *θ*_1_, or *φ*_1_ are then plotted.

Furthermore, Type I error and power for each DGP and each statistical method are calculated by estimating the Type I error by setting the parameter of interest equal to (**DGP1** : *ρ*_0_ = *ρ*_1_ = 0, **DGP2** : *θ*_0_ = *θ*_1_ = 3, **DGP3** : *φ*_0_ = *φ*_1_ = 3). To assess power, a reference point for group 0 (*ρ*_0_ = *φ*_0_ = 0, *θ*_0_ = 3) is set, while setting the parameter of interest of group 1 to values surrounding group 0’s (*ρ*_1_ = *φ*_1_ = (−.4, −.2, .2, .4), *θ*_1_ = (1, 2, 3, 4)). The Type I error and power are evaluated in a range of sample sizes (*n*_0_ = *n*_1_ = 500, 1000, 1500) and a fixed number of predictors ((p = 10)) to evaluate changes in the estimator as sample sizes increase.

#### 2.2.10. Inflammatory biomarkers and effect on dementia

Finally, the effect that inflammatory biomarkers collectively had on dementia risk is analyzed by performing the three aforementioned methods on the partial covariance structure of non-dementia and dementia group.

## 3. Results

### 3.1. Simulation Analysis

This section summarizes the results of the simulation analysis as described in Section 2.2.9.

#### 3.1.1. Test Statistics and P-values Results

Figure 2 depicts the graphs of the three test statistics and the respective p-values (Figure 3) for the three different simulations. Note that the parameters for **DGP2** (*θ*_1_) are scaled to match the other simulations in the overlying illustration. As a reminder, in the original simulation (*θ*_1_ ∈ (1, 5), *θ*_0_ = 3), this is re-scaled to 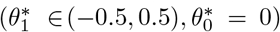 in graphical presentation. The test statistics in **DGP1** do not change noticeably with the parameters, although their corresponding p-values changed in a predictable way with the exception of **DGP2**. The Tracy-Widom distribution performed well under **DGP1** and **DGP3** as it exhibits symmetry in the test statistic and the p-value graph. However, the **DGP2** simulation was not symmetric but was more significant on the right side. The Tracy-Widom p-value did not reach significance level (*α* = .05) even at 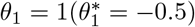, the lowest value in **DGP2** simulation. In fact, the minimum of the Tracy-Widom test statistic (the maximum p-value) was reached when *θ*_1_ *<* 3, lower than the true null case. Lastly, the scaled Parametric Bootstrap statistic had good results in all three simulation methods.

**Figure 2:**
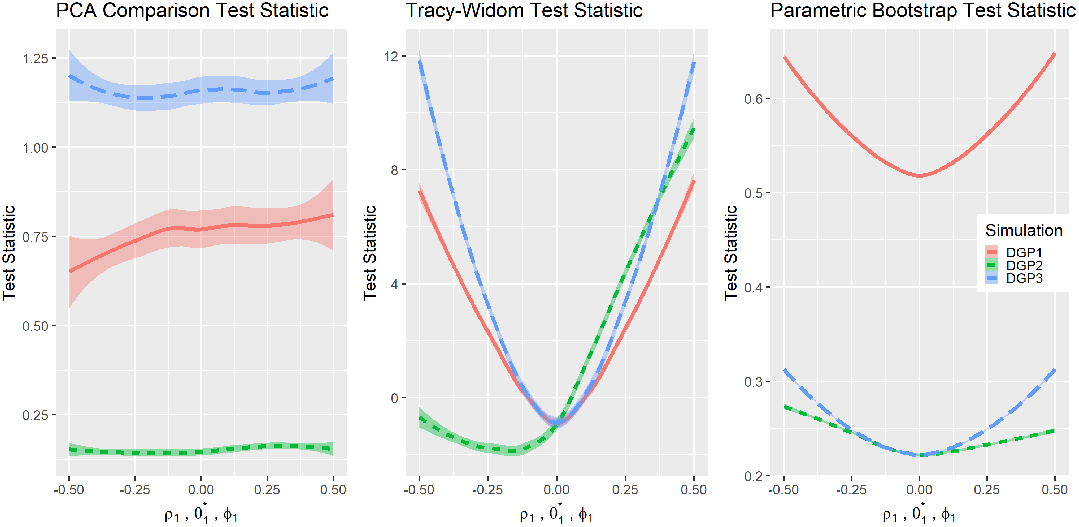
Comparisons of test statistics

**Figure 3:**
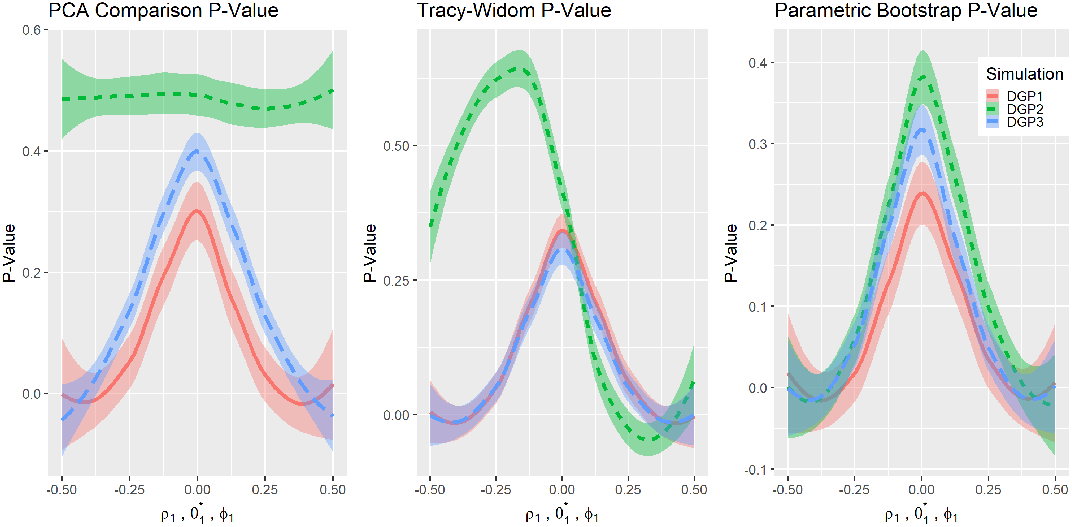
P-value comparisons

The one-sided peculiarity of Tracy-Widom’s distribution in **DGP2** is worth exploring. A compelling argument for this phenomenon is that the size or determinant of Σ_1_ has an one-sided effect on the Tracy-Widom distribution, as *S*_1_ is used in the numerator to make the combined covariance matrix. Indeed, a simulation that varies the matrix size clearly shows this effect, we call this simulation **DGP4**.

##### DGP4

Data was generated from independent bivariate normal distributions with zero-mean and variance 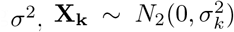. Where *k* = (0, 1) and 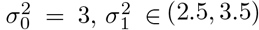.

**DGP4** simulation result (Figure 4) clearly showed the one-sided relationship between the Tracy-Widom statistic and the difference between the variances 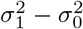. Therefore, the appropriate way to test the equality of the covariance matrices would be by means of a two-sided p-value. However, this contradicts the results of **DGP1** and **DGP3**, which showed that the Tracy-Widom statistic is two-sided with respect to the difference in covariance between variables. This contradiction creates a difficult dilemma. On the one hand, there is the option of looking for differences in magnitude in variances, where a one-sided p-value would be appropriate. On the other hand, if the objective is to determine differences in correlation between variables, the two-sided p-value would be the appropriate choice. For sake of simplicity, the data analysis focuses on the latter by standardizing the variables of interest. The standardization to zero mean and unit variance is consistent with similar analyses, such as in (Frost et al., 2016).

**Figure 4:**
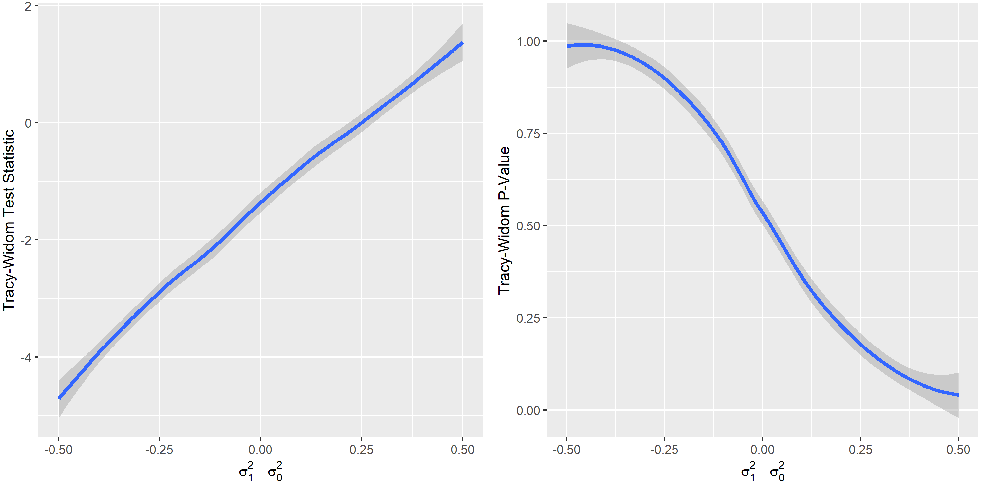
DGP4 Simulation Results

#### 3.1.2. Type I Error and Power Results

Next, the three methods using Type I error and power of the tests were assessed. Type I error is represented in the columns where the parameters for both groups are equal to (*ρ*_1_ = 0, *θ*_1_ = 3, and *φ*_1_ = 0,). Although the PCGC method captures the true Type I error (*α* = .05) quite well, the power is relatively low in all simulations; in fact, the power does not appear to change at all in **DGP2**. The Parametric Bootstrap method accurately estimates Type I error, while also having high power for all simulations. Meanwhile, the Tracy-Widom method was consistently conservative in estimating the Type I error in all simulations, with the **DGP1** simulation being the most conservative. As for power, the Tracy-Widom method had good power distribution in **DGP1** and **DGP3**, although is in general less powerful than the Parametric Bootstrap method. However, the Tracy-Widom method was extremely under-powered in **DGP2** when *θ*_1_ *< θ*_0_, this phenomenon is a consequence of Tracy-Widom method’s non-symmetric interaction with **DGP2**.

In general, power increases as the sample size increases, whereas the Type I error does not have any discernible changes. Additionally, the power increases along with sample sizes given a fixed number of variables; this is true for all three methods.

In summary, the Parametric Bootstrap method had the most impressive simulation results; not only did it reasonably estimate Type I error, but it also demonstrated high power in all the simulations. The least impressive method was PCGC. While it had reasonable Type I power estimation, it was under-powered in comparison. In addition, PCGC was completely unable to detect the parameter changes in **DGP2**. The Tracy-Widom method was conservative in estimating the Type I error in all simulations. It had high power in **DGP1** and **DGP3**, although not as much as the Parametric Bootstrap. However, the Tracy-Widom method was extremely underpowered in **DGP2** when (*θ*_1_ *< θ*_0_) as mentioned previously.

### 3.2. Analysis of Inflammatory Biomarker Data

This section summarizes the results of the analysis of the data on the FHS inflammatory biomarkers and dementia.

#### 3.2.1. Preliminary Analysis

These analyzes sought evidence suggesting structural differences in biomarkers between participants in the dementia and non-dementia group. Figure 5 illustrates the comparison between the partial correlation of the two groups using a heat map visualization. There seems to be some negative correlation among some variables in the dementia group, which does not appear in the non-dementia group.

**Figure 5:**
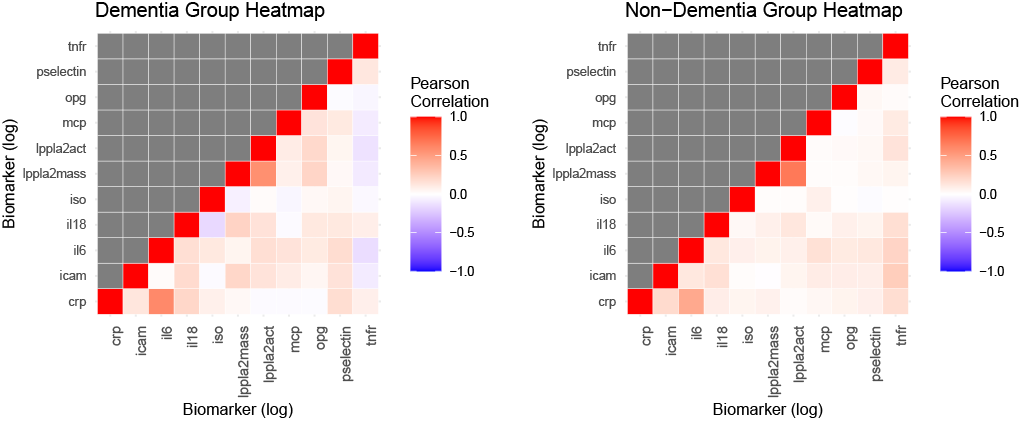
Heatmap Comparison

Another possible way to visualize the structural differences in the partial covariance matrices between the non-dementia group and the dementia group is by comparison of the scree plots (Figure 6). In both groups the eigenvalues of the first two components appear to be very similar, while there exists minor differences in the latter components.

**Figure 6:**
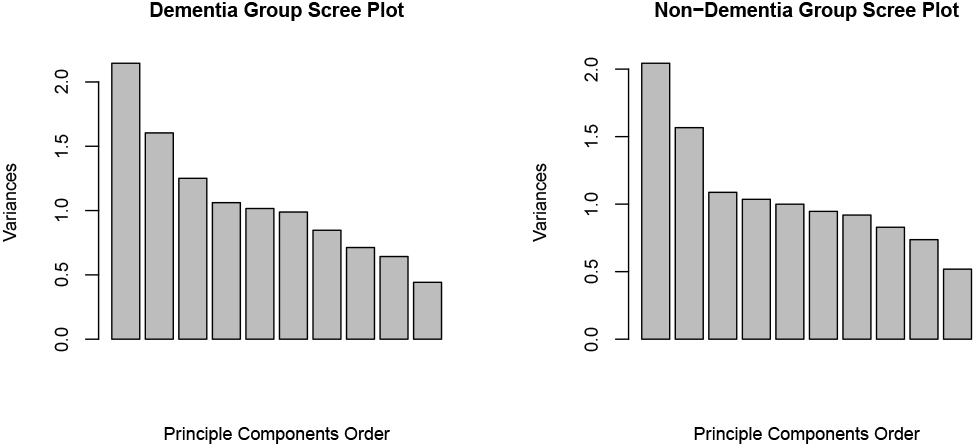
Scree-plot comparison

#### 3.2.2. Results of the Inferential Procedures

The previous sections considered inflammatory biomarkers and the difference in the structure of the covariance between the two groups by generating a simple covariance matrix and the partial covariance matrix. The simple covariance matrix was calculated directly from the log-transformed biomarkers, while the partial covariance was calculated from the residuals after adjusting for the covariates. The results from analyzing the simple covariance would show whether there is a difference in covariance structure by itself, while the results from the partial covariance would show whether differences exist after adjusting for confounding factors.

The results show significant differences between the covariance matrices in all cases except for the PCGC method on the simple covariance matrix. Given the lackluster performance of PCGC in the simulations, the results of Tracy-Widom and Parametric Bootstrap seem more reasonable. Therefore, it appears that there is a significant difference between the simple covariance and partial covariance of the log-adjusted biomarkers of the dementia group and the non-dementia groups. Partial covariance appears to be more significant than simple covariance, which gives credence to covariate adjustments. This result explicitly reveals differences in the dementia groups that the preliminary visualizations did not clearly show.

## 4. Discussion

Despite extensive research on individual risk factors for dementia, there are far fewer studies that consider how these risk factors synergistically influences dementia. Current solutions to multivariate problems (such as machine learning) are complex given the large and varied data involved. On the other hand, statistical methods based on comparison of covariance matrices have been applied to a variety of problems, but not to biological risk factors of dementia. The purpose of this study was to test simpler covariance-based models that summarize multiple variables into a single test and score by efficiently examining the combined effect of many variables.

Three methods were investigated that combined the covariance matrices of two groups and then were summarized as a test statistic to test whether the matrices are structurally different. The validity of these methods via simulation, Type I error, and power analysis suggests that the Parametric Bootstrap method is effective in identifying both significant and insignificant differences between covariance matrices. By comparison, the PCGC method has low power and was unreliable in some cases. The Tracy-Widom method was conservative in estimating the Type I error, but did accurately estimate power. However, for non-normal distributed data, there were conflicting effects of variable variances and inter-variable covariances using the Tracy-Widom Statistics. These results indicate that the Parametric Bootstrap and Tracy-Widom methods were reliable methods for analyzing approximately normally distributed data.

The application of these methods to the clinical question of inflammatory biomarkers and their relationship with dementia risk found that these tests detected significant differences in the covariance structure of biomarkers between the dementia group and the non-dementia group. This result reinforces the established consensus that inflammation biomarkers are indeed risk factors for dementia. Furthermore, the 11 biomarkers analyzed in this paper can act as a collective risk factor for dementia. The collective covariance effect is certainly significant for dementia regardless of the significance of the individual mean effects. There are limitations both in the methods and with the data. These methods are not well tested and thus are not established theoretically, which leads to a reliance on the use of simulation to assess the validity of these methods. Although these methods did determine the existence of a significant difference between two groups collectively among biomarkers, they could not determine which variables are different. Furthermore, the data was not col-lected from the same period. Interleukin-18 (Il18) samples were measured from blood from the 7^*th*^ health exam (1998 - 2001), while the rest were measured from health exam 8 (2005 - 2008).

Furthermore, the covariate information came mainly from exam 8 values; exam 7 values were used only when exam 8 values are unavailable. These compromises could introduce additional noise that could not be controlled in order to maximize the number of biomarkers in the study. Furthermore, there was an imbalance in sample sizes between the non-dementia group (n = 2536) and the dementia group (n = 168). Imbalanced data introduces bias (King and Zeng, 2001) and ‘wastes’ the extra samples in the non-dementia group. Future research is warranted to determine theoretical justifications for the parametric Bootstrap and Tracy-Widom methods using more balanced study groups.

In summary, Parametric Bootstrap and Tracy-Widom methods are valid candidates for testing the significance of covariance differences among standardized, multivariate variables between groups, while the PCGC method is unreliable. Parametric Bootstrap appears to have suitable Type I error estimates and strong power, while Tracy-Widom has conservative Type I error estimates but good power. In a clinical research application, data analysis showed significant differences in protein biomarker covariance matrices between the non-dementia group and the dementia group. This result is expected, given the literature on inflammatory biomarkers as a risk factor for dementia. Utilizing new and cutting-edge analytic approaches can further enrich the understanding of inflammatory biomarkers and their effects on dementia.

## 5. Acknowledgments

This paper is generously supported by Framingham Heart Study’s National Heart, Lung, and Blood Institute contract (N01-HC-25195), and the National Institute on Aging (AG016495, AG008122, AG062109, AG068753)

## Footnotes

1 PCGC - Principal Components Group Comparison, PBoot - Parametric Bootstrap, TW - Tracy-Widom

